# DJ-1 deficiency and aging: dual drivers of retinal mitochondrial dysfunction

**DOI:** 10.1101/2025.05.19.654941

**Authors:** Mala Upadhyay, Johnathon Sturgis, Sanghamitra Bhattacharyya, Ke Jiang, Stephanie A. Hagstrom, Vera L Bonilha

**Author notes:** Correspondence: Vera L. Bonilha, Cleveland Clinic Lerner College of Medicine, The Cole Eye Institute, i31, 9500 Euclid Avenue, Cleveland, OH, 44195, USA.

## Abstract

We have previously extensively characterized the role of DJ-1 in oxidative stress regulation in the retina and RPE during aging. However, the DJ-1 protein also plays a role in regulating mitochondria’s response to oxidative stress by translocating to the mitochondria where it helps clear generated reactive oxygen species (ROS). To study the effects of aging and oxidative stress in the retina, the DJ-1 KO mouse was analyzed. Freshly dissected ex vivo retinal punches were analyzed for real-time live cell metabolism. Total DNA and protein were isolated from RPE, and retina of 3- and 15-month-old DJ-1 WT and DJ-1 KO mice. The mitochondrial DNA (mtDNA) genome was divided into four discrete regions (RI–RIV), and lesions/10kb were quantified using long-extension PCR. mtDNA content was analyzed using RT-qPCR. Protein levels of OXPHOS complexes, POLG, OGG1, SOD2, and PGC1α were measured by western blotting. Seahorse analysis detected significantly decreased basal and maximal OCR in 3- and 15-month-old DJ-1 KO compared to age-matched DJ-1 WT. In the RPE, a significant decrease in the protein levels of the NDUFB8 subunit of CI, the SDHB subunit of CII, and MTCO1 of CIV in 15-month-old DJ-1 KO mice compared to 15-month-old DJ-1 WT, while the ATP5A subunit of CV was significantly decreased in 3-month-old DJ-1 KO mice compared to 3-month-old DJ-1 WT. In the retina, significantly decreased levels of NDUFB8 subunit of CI and MTCO1 of CIV were detected in the in 3-month-old DJ-1 KO mice compared to 3-month-old DJ-1 WT. We observed a significant increase in mtDNA gene content in 15-month-old DJ-1 KO RPE and retina compared to age-matched DJ-1 WT. The PGC1α levels significantly decreased in 3- and 15-month-old RPE lysates compared to their age group DJ-1 WT. However, in the retina, there was only a decrease in DJ-1 WT with aging. The POLG levels increased when 15-month-old DJ-1 KO lysates were compared to 15-month-old DJ-1 WT. The mtDNA lesions in the RPE detected a trend of increase in 15-month-old DJ-1 WT and DJ-1 KO RPE in all regions compared to 3-month-old mice. In the retina, a significant increase in mtDNA lesions/10kb accumulation in the 15-month-old DJ-1 WT RIV was detected compared to 3-month-old DJ-1 WT. The OGG1 levels significantly increased in 3-month-old retinal lysates compared to the 3-month-old DJ-1 WT. Our findings suggest that DJ-1 is critical for mitochondrial regulation and function in RPE and retina. The observed changes reflect mitochondrial dysfunction related to absence of DJ-1.

## 1. INTRODUCTION

Age-related macular degeneration (AMD) is characterized by a progressive loss of the cone-rich macula, the central region of the retina needed for high visual acuity and color vision, and the surrounding vasculature (1). AMD primarily affects the outer retina, which includes the RPE, Bruch membrane, the choriocapillaris, and the underlying choroid. Genetic factors, smoking, inflammation, oxidative stress, and, above all, aging, have been suggested as primary factors associated with the pathogenesis of AMD (2–5). The exact mechanism of AMD pathogenesis is still a mystery. However, multiple studies have implicated mitochondrial dysfunction as a promoting factor for AMD pathogenesis (6–10).

Mitochondria are double-membrane organelles engaged in adenosine triphosphate (ATP) production through oxidative phosphorylation (OXPHOS). Mitochondria contribute to many processes central to cellular function and dysfunction, including calcium signaling, cell growth and differentiation, cell cycle control, and cell death (11, 12). Mitochondria are also unique organelles because they possess their circular, double-stranded DNA, are intron-less, separate from the nucleus, and replicate independently of the cell cycle, irrespective of the mitotic phase (13). The mammalian mitochondrial DNA (mtDNA) encodes for 13 OXPHOS complex proteins, two rRNAs, and 22 tRNAs. mtDNA-encoded proteins and RNAs are involved in mtDNA gene expression and, therefore, are critical for driving the OXPHOS system (14). Each eukaryotic cell contains multiple maternally inherited copies of mtDNA, and their copy number varies with tissue type, developmental stage, and metabolic needs (15). Mutations due to replication error or incorrect base pairing have been identified in its tRNAs, rRNAs, and protein-coding genes, compromising gene expression and causing OXPHOS deficiency. The mutations in multicopy mtDNA can be homoplasmic or heteroplasmic; heteroplasmic mutation attaining a certain threshold being associated with common age-associated human disease (16–18). Mitochondria generate 90% of ATP but also generate reactive oxygen species (ROS) due to electron leakage by complex I (C I) and III (C III) of the OXPHOS system (19). At a steady level, less reactive mitochondrial ROS (mt ROS) in the form of H2O2 participate in cellular signaling and maintain homeostasis (20). With aging, there is a cumulative accumulation of highly reactive mtROS, leading to mitochondrial dysfunction; this, in turn, becomes a predisposition factor for age-related diseases (21).

DJ-1, also known as PARK7, is a protein renowned for its ability to protect neurons from oxidative stress (22–24). The DJ-1 protein plays a pivotal role in regulating mitochondria’s response to oxidative stress by translocating to the mitochondria, which aids in the clearance of generated ROS (25, 26). In conditions of oxidative stress, DJ-1 has been shown to localize to mitochondria (26, 27) and maintain mitochondrial C I activity (28). This robust defense mechanism underscores the potential of DJ-1 in combating oxidative stress, a key factor in the pathogenesis of AMD. Indeed, our previous research has reported an increased presence of DJ-1 in isolated human RPE and choroid/BM and, comparatively, more DJ-1 localization in AMD donor eyes vs control non-AMD eyes (29). We also detected increased levels of oxidized DJ-1, an important finding as over oxidizing of DJ-1 results in inactivation of the protein.

Our research has shown that the absence of DJ-1 leads to progressive RPE/retinal degeneration with increased RPE/retinal oxidative stress, suggesting a protective role of DJ-1 in the retina with aging (30, 31). More recently, we extensively characterized the role of DJ-1 in regulating oxidative stress in the retina and RPE during aging (32). Building on our previous work, the present study we undertook a comprehensive analysis of the mitochondrial function in RPE and retina during aging using young (3-month-old) and aged (15-month-old) DJ-1 knockout (KO) and wild-type (DJ-1 WT) mice. Our findings suggest that loss of DJ-1 contributes to mitochondrial dysfunction by modulating OXPHOS subunit expression, mitochondrial DNA content, and mitochondrial biogenesis. These findings not only enhance our understanding of the role of DJ-1 in mitochondrial function in the retina and RPE but also provide potential targets for therapeutic interventions in AMD and other age-related retinal diseases.

## 2. RESULTS

### 2.1 Loss of DJ-1 lowers mitochondrial oxygen consumption in the retina

Our previous unbiased proteomics analysis of retinal and RPE lysates from 3-month-old mice identified increased levels of mitochondrial proteins in the retinas and decreased levels of mitochondrial proteins in the RPE of the DJ-1 KO mice compared to control mice (32). To further determine how DJ-1 deficiency and aging impacted the retinal metabolism, we conducted a Seahorse analysis on freshly dissected ex vivo retinal tissues, measuring the two major energy-producing pathways in the cell, namely oxidative phosphorylation (OXPHOS) and glycolysis. We first quantified the oxygen consumption rate (OCR) on 3- and 15-month-old DJ-1 WT and DJ-1 KO 1 mm retinal punches using a previously described method to access OXPHOS respiration levels (33). The OCR was measured at baseline and after each drug injection (Fig. 1A). Basal OCR was significantly reduced in the retina of 3-month-old DJ-1 KO compared to 3-month-old DJ-1 WT mice. A similar significant reduction was observed in the 15-month-old DJ-1 KO mice’s retina compared to 15-month-old DJ-1 WT (Fig. 1B). We did not observe any significant changes in retinal OCR with aging when 15-month-old DJ-1 KO were compared to 3-month-old DJ-1 KO mice and 15-month-old WT mice were compared to 3-month-old WT mice. Following the injection of uncoupler Bam15, an agent that dissipates the proton gradient, oxygen consumption rises and reaches a maximum level. Like basal respiration, maximal OCR was significantly reduced at baseline in the retina of the 3-month-old DJ-1 KO compared to age-matched DJ-1 WT mice. A similar significant decrease was observed at 15 months of age when comparing DJ-1 KO mice to DJ-1 WT mice. No significant changes in maximal OCR were observed with aging when comparing 15-month-old mice to 3-month-old in both DJ-1 WT and DJ-1 KO groups (Fig. 1C). Oxygen consumption drops to bottom level following the injection of OXPHOS complex inhibitors rotenone and antimycin A. Mitochondrial reserve capacity (MRC) was calculated for each age and genotype group. MRC was significantly higher in 3-month DJ-1 KO compared to 3-month DJ-1 WT mice (Fig. 1D). No significant differences in MRC were observed with aging, nor between 15-month-old DJ-1 KO and 15-month-old DJ-1 WT mice.

**Figure 1.**
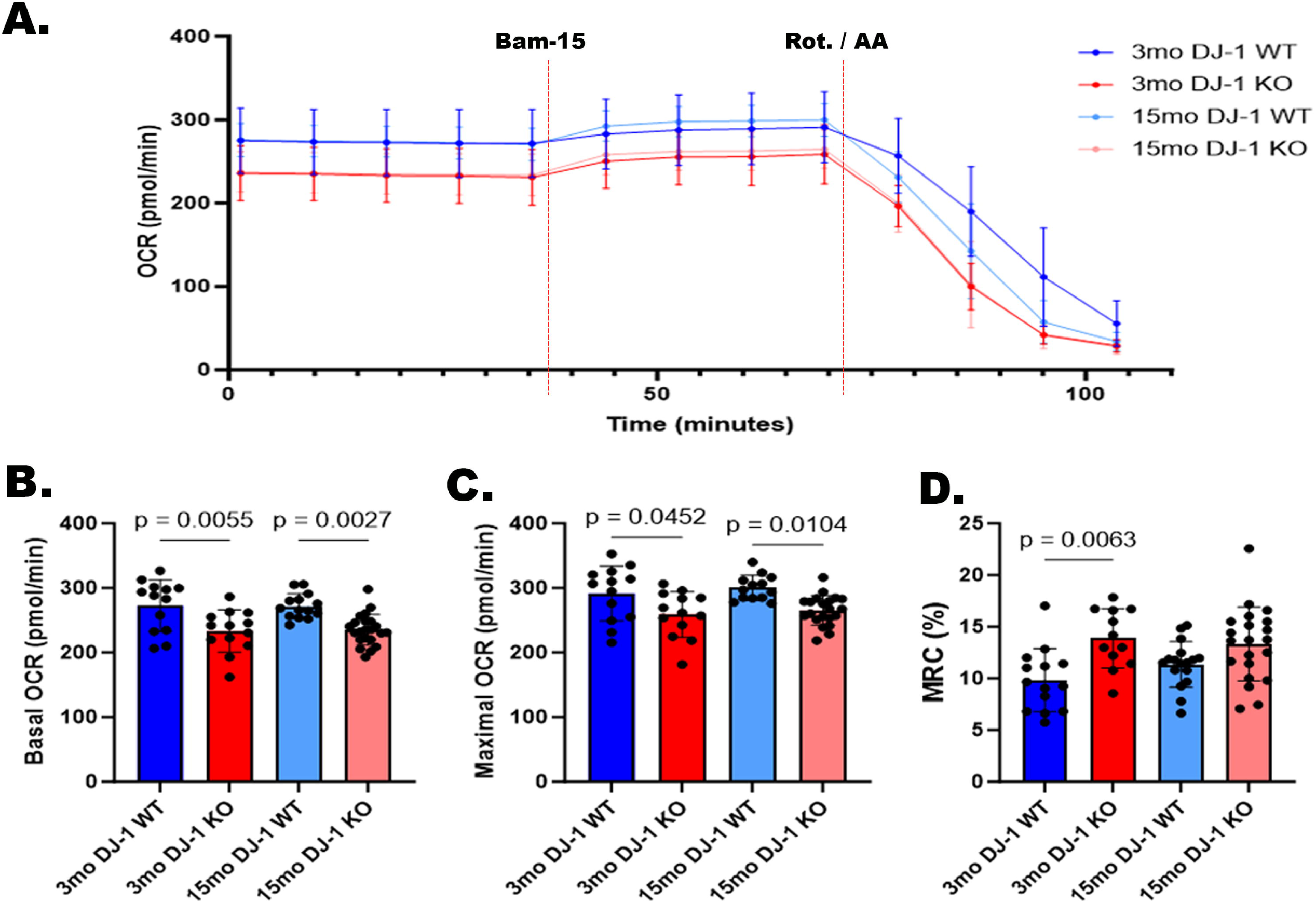
Effects of aging and DJ-1 loss in mitochondrial bioenergetics of retinas by Seahorse Analyzer. Five measurements were taken before treatment (basal respiration), four measurements were taken after 5 μM Bam15 injection (maximal respiration), and four measurements were taken after 1 μM rotenone/antimycin A injection (mitochondrial reserve capacity). A) Unnormalized OCR measurements in retinal punches from 3-month-old DJ-1 WT (dark blue), 3-month-old DJ-1 KO (dark red), 15-month-old DJ-1 WT (light blue), and 15-month-old DJ-1 KO (light red) mice. B) Basal OCR, C) Maximal OCR, and D) Mitochondrial reserve capacity (MRC) of mice described above. Error bar indicates SEM. Data points = technical replicates / individual retina punches from n = 3-5 mice. Statistics: one-way ANOVA, significance: p≤0.05 depicted in the figure.

Next, we quantified glycolysis using the Seahorse analyzer by measuring the extracellular acidification rate (ECAR) to determine the impact of DJ-1 deficiency on the retinal glycolytic flux. The ECAR of retinal punches was measured at baseline, after injection of OXPHOS inhibitors rotenone and antimycin A, and after injection of a glucose analog 2-Deoxy-D-glucose (2-DG) (Suppl. Fig. 1A). Basal ECAR was significantly reduced at baseline in the retina of 3-month-old DJ-1 KO compared to 3-month-old DJ-1 WT mice. No significant changes in basal ECAR were observed between the retinas of 15-month-old DJ-1 KO and 15-month-old DJ-1 WT mice. Moreover, no significant changes in ECAR were observed in the retina with aging in 15-month-old DJ-1 WT compared to 3-month-old DJ-1 WT mice (Suppl. Fig. S1B). Following the injection of rotenone and antimycin A, maximal ECAR can be observed due to the increase in glycolytic flux being driven by inhibition of the electron transport chain. Maximal ECAR showed no significant difference between genotypes and with aging (Suppl. Fig. 1C). After injection of the glucose analog 2-DG, glycolysis is inhibited, and ECAR is significantly reduced. Similarly, we calculated glycolytic reserve capacity (GRC). The GRC was significantly higher in the retina of 15-month DJ-1 WT and DJ-1 KO retinas when compared to the 3-month-old DJ-1 WT and DJ-1 KO mice, respectively, but showed no differences when retinas of the same age were compared (Suppl. Fig. 1D). Altogether, these observations suggested DJ-1 loss might lead to energy deficiency driven by reduced basal mitochondrial respiration and glycolysis in retina that can be detected at 3-months of age. Although aging within the genotypes, did not change the basal OCR outcome, a significant basal and maximal OCR reduction at the 15-month timepoint between the two genotypes suggest reduced OXPHOS as a measure of mitochondrial dysfunction remains a factor with age.

### 2.2 Loss of DJ-1 alters OXPHOS subunit expression in RPE and retina

Previous studies detected the interaction and influence of DJ-1 in the assembly of CI, CIV, and CV in different cells (34). To determine if loss of DJ-1 can alter mitochondrial function by affecting the OXPHOS system, we incubated the resolved proteins from 3- and 15-month-old DJ-1 WT and DJ-1 KO RPE lysates with a total OXPHOS cocktail antibody, which detects one subunit of each of the five OXPHOS complexes (Fig. 2A). Western blot analysis of RPE lysates from 3-month-old and 15-month-old DJ-1 WT and DJ-1 KO mice detected similar protein levels of the NDUFB8 subunit of CI and the SDHB subunit of CII in 3-month-old DJ-1 KO and DJ-1 WT. With aging, levels of NDUFB8 and SDHB remained comparable to young DJ-1 WT and DJ-1 KO, but there was a significant decrease in its expression in 15-month-old DJ-1 KO mice compared to 15-month-old DJ-1 WT, suggesting a role of DJ-1 in sustaining NDUFB8 and SDHB expression in RPE (Figs. 2B, 2C). No significant changes in the levels of the UQCRC2 subunit of CIII between DJ-1 WT and DJ-1 KO mice and with aging were detected (Fig. 2D). Protein levels of the MTCO1 (cytochrome c oxidase subunit I) of CIV were again comparable between 3-month-old DJ-1 KO and DJ-1 WT and significantly decreased between 15-month-old DJ-1 WT and 15-month-old DJ-1 KO (Fig. 2E). Notably, the protein levels of the ATP5A subunit of CV were significantly decreased in 3-month-old DJ-1 KO mice compared to 3-month-old DJ-1 WT, and these levels were comparable to both 15-month-olf DJ-1 WT and DJ-1 KO RPE (Fig. 2F).

**Figure 2.**
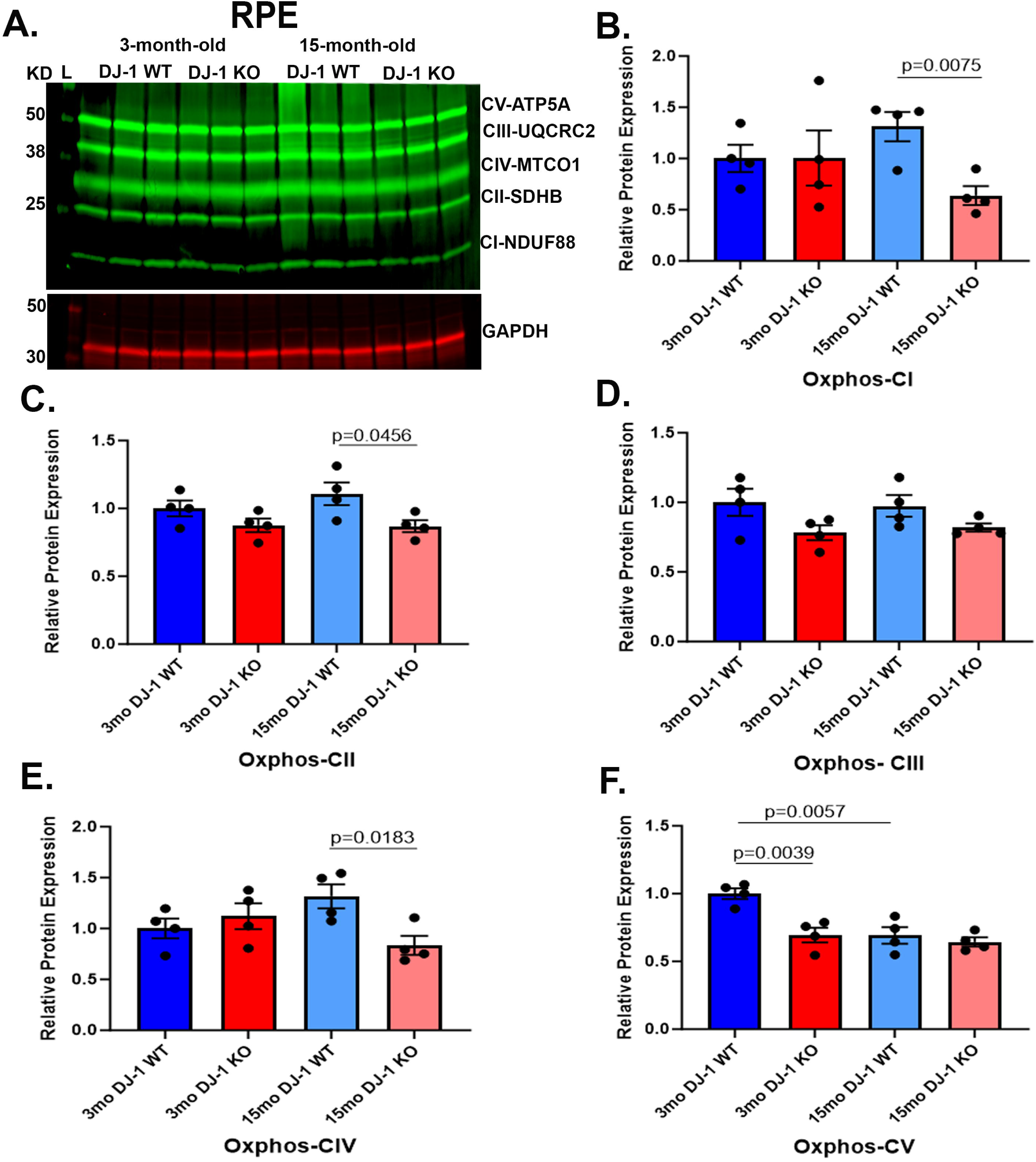
Effects of aging and DJ-1 loss on levels of OXPHOS in RPE. A) Representative western blot of OXPHOS subunits and GAPDH in RPE lysate from 3- and 15-month-old DJ-1 WT and DJ-1 KO mice. B-F) Quantitative analysis and graphic representation of NDUFB8 (CI), SDHB (CII), UQCRC2 (CIII), MTCO1 (CIV), and ATP5A (CV) signal intensity in RPE lysate from 3-month-old DJ-1 WT (dark blue bar), 3-month-old DJ-1 KO (dark red bar), 15-month-old DJ-1 WT (light blue bar), and 15-month-old DJ-1 KO (light red bar) mice. GAPDH was used as a housekeeping normalizer, and relative protein expression was calculated using 3-month-old DJ-1 WT as a normalizer. Data are represented as mean ± SEM; n=3-4 mice per group; statistical analysis using unpaired Student’s t test; significance: p≤0.05 depicted in the figure.

To determine if the loss of DJ-1 can alter mitochondrial function by affecting the OXPHOS system, we also incubated resolved proteins from 3- and 15-month-old DJ-1 WT and DJ-1 KO retina lysates with the total OXPHOS cocktail antibody described above (Fig. 3A). Western blot analysis of retina lysates from 3-month-old and 15-month-old DJ-1 WT and DJ-1 KO mice detected significantly decreased levels of NDUFB8 subunit of CI in both 3- and 15-month-old DJ-1 KO compared to DJ-1 WT (Fig. 3B), suggesting a role of DJ-1 in sustaining NDUFB8 expression in RPE. No significant changes were detected in the levels of the SDHB subunit of CII and the UQCR2 subunit of CIII between DJ-1 WT and DJ-1 KO in both 3- and 15-month-old mice (Fig. 3C, 3D). Protein levels of the MTCO1 of CIV were significantly decreased between 3-month-old DJ-1 WT and DJ-1 KO retinas and between 3- and 15-month-old DJ-1 WT (Fig. 3E). The protein levels of the ATP5A subunit of CV were not significantly altered between DJ-1 WT and DJ-1 KO in 3- and 15-month-old mice (Fig. 3F). Our data suggested that DJ-1 deficiency can negatively alter OXPHOS subunit expression in RPE and retina at baseline levels and with aging. However, different subunits were affected in the RPE and retina.

**Figure 3.**
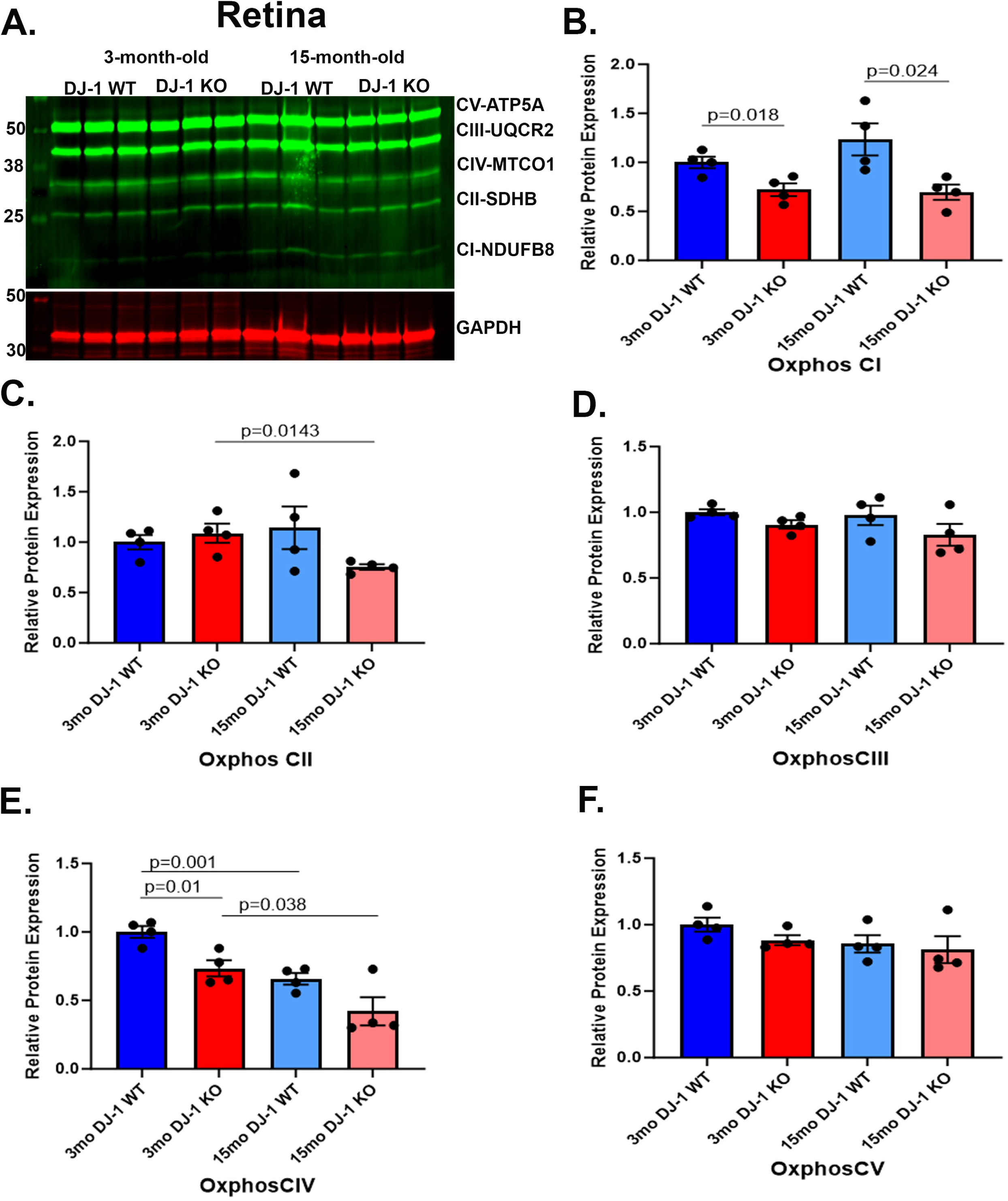
Effects of aging and DJ-1 loss on OXPHOS levels in the retina. A) Representative western blot of OXPHOS subunits and GAPDH in retina lysate from 3- and 15-month-old DJ-1 WT and DJ-1 KO mice. B-F) Quantitative analysis and graphic representation of NDUFB8 (CI), SDHB (CII), UQCRC2 (CIII), MTCO1 (CIV), and ATP5A (CV) signal intensity in RPE lysate from 3-month-old DJ-1 WT (dark blue bar), 3-month-old DJ-1 KO (dark red bar), 15-month-old DJ-1 WT (light blue bar), and 15-month-old DJ-1 KO (light red bar) mice. GAPDH was used as a housekeeping normalizer, and relative protein expression was calculated using 3-month-old DJ-1 WT as a normalizer. Data are represented as mean ± SEM; n=3-4 mice per group; statistical analysis using unpaired Student’s t test; significance: p≤0.05 depicted in the figure.

### 2.3 Loss of DJ-1 modulates mitochondrial DNA content in RPE and retina

Previous studies determined that mitochondrial DNA (mtDNA) copy number is a biomarker of mitochondrial function and overall health and that levels of mtDNA copy number are associated with several age-related diseases (35–38). To further determine how DJ-1 deficiency and aging impacted mitochondrial health, we analyzed the relative mtDNA content of the mitochondrial genes 16S rRNA, a ribosomal RNA molecule found in the 30S subunit of prokaryotic ribosomes, and mitochondrially encoded cytochrome c oxidase I (mtco1) in the RPE of 3- and 15-month-old DJ-1 KO and DJ-1 WT mice. The relative mtDNA content was almost a two-fold increase in both mtDNA genes in 3-month-old DJ-1 KO mice compared to DJ-1 WT RPE. Furthermore, there was a three-fold increase in both mtDNA gene content in 15-month-old DJ-1 KO mice compared to 15-month-old DJ-1 WT RPE (Figs. 4A, 4B). In the face of these results, we proceeded to evaluate the levels of peroxisome proliferator-activated receptor gamma coactivator 1 alpha (PGC1α), a coactivator transcription factor known as the primary driver of mitochondrial biogenesis in most tissues (39). Western blot analysis of RPE lysates in both 3- and 15-month-old DJ-1 KO compared to DJ-1 WT detected significantly decreased levels of PGC1α. The levels of PGC1α were also significantly decreased in the RPE of DJ-1 WT with aging (Figs. 4C, 4D). We also analyzed the levels of the exonuclease domain of the polymerase gamma (POLG) protein, which is responsible exclusively for mtDNA replication and repair (40). The RPE POLG levels were significantly decreased with aging in the DJ-1 KO (Figs. 4E, 4F).

**Figure 4.**
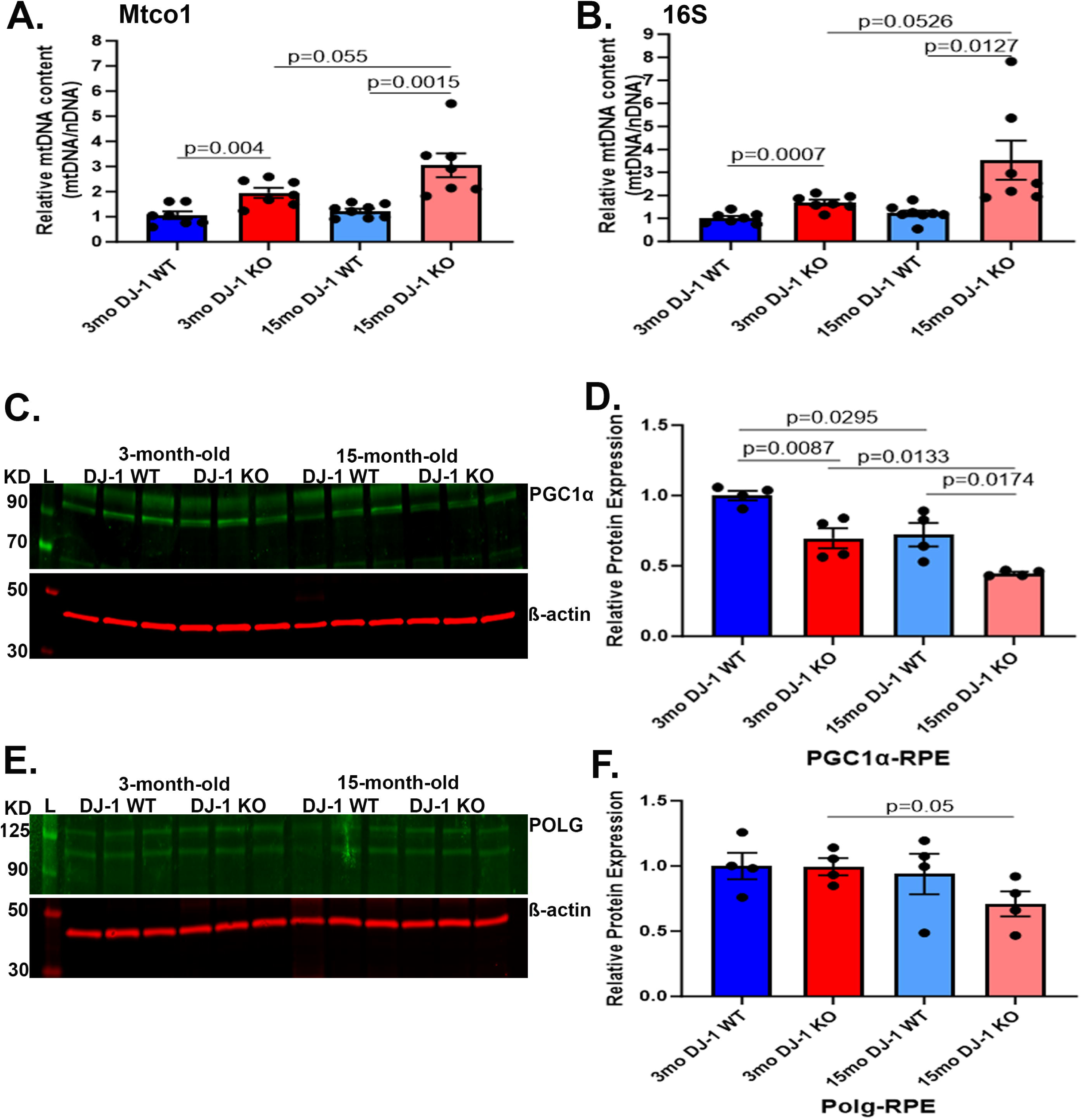
Effects of aging and DJ-1 loss in mtDNA content and mitochondrial biogenesis in the RPE. A-B) Real-time PCR quantitative analysis of Mtco1 and 16S mtDNA content normalized to nuclear β-globin in 3-month-old DJ-1 WT (dark blue bar), 3-month-old DJ-1 KO (dark red bar), 15-month-old DJ-1 WT (light blue bar), and 15-month-old DJ-1 KO (light red bar) RPE. C) Representative western blot for PGC1α and β-actin in RPE lysates from 3- and 15-month-old DJ-1 WT and DJ-1 KO mice. D) Quantitative analysis and graphic representation of PGC1α signal intensity in RPE from lysates described above. E) Representative western blot for POLG and β-actin from 3- and 15-month-old DJ-1 WT and DJ-1 KO mice. F) Quantitative analysis and graphic representation of POLG signal intensity in RPE lysates described above. β-actin was used as a housekeeping normalizer, and relative protein expression was calculated by using 3-month-old WT as a normalizer. Data are represented as mean ± SEM; n=3-4 mice per group; statistical analysis using unpaired Student’s t test; significance: p≤0.05 depicted in the figure.

A similar analysis was performed in retina lysates (Fig. 5). The relative mtDNA content in the retina was comparable between 3-month-old DJ-1 WT and DJ-1 KO. With aging, a significant increase in relative mtDNA content was observed in 15-month-old DJ-1 KO mice compared to 15-month-old DJ-1 WT (Figs. 5A, 5B). Western blot analyses of PGC1α in the retina detected a decreasing trend in 3-month-old DJ-1 KO compared to 3-month-old DJ-1 WT mice. Interestingly, the expression of PGC1α protein significantly decreased in 15-month-old DJ-1 WT compared to 3-month-old DJ-1 WT (Figs. 5C, 5D). We also analyzed POLG expression in all the groups. The levels of POLG were comparable between 3-month-old DJ-1 WT and DJ-1 KO mice. With aging, the 15-month-old DJ-1 KO displayed significantly increased levels of POLG compared to 15-month-old DJ-1 WT (Figs. 5E, 5F). These observations suggested that DJ-1 is critical for increasing mtDNA content in RPE and that these changes were inversely related to PGC1α. In the retina, DJ-1 loss elicited increased mtDNA content in the retinas of DJ-1 KO mice, concomitant with no significant changes in the levels of PGC1α.

**Figure 5.**
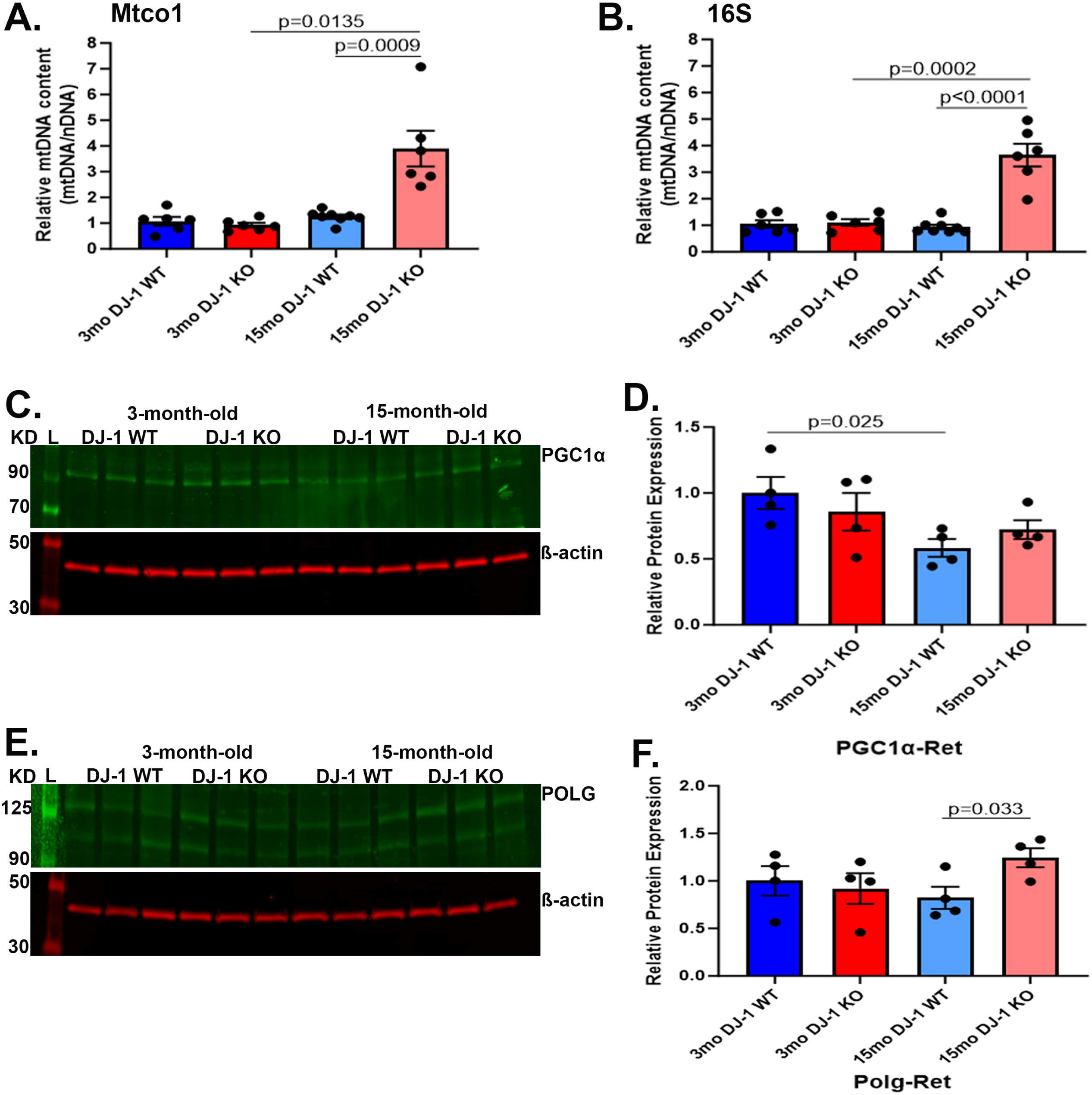
Effects of aging and DJ-1 loss on the retina’s mtDNA content and mitochondrial biogenesis. Real-time PCR quantitative analysis of Mtco1 and 16S mtDNA content normalized to nuclear β-globin in 3-month-old DJ-1 WT (dark blue bar), 3-month-old DJ-1 KO (dark red bar), 15-month-old DJ-1 WT (light blue bar), and 15-month-old DJ-1 KO (light red bar) RPE. C) Representative western blot for PGC1α and β-actin in retina lysate from 3- and 15-month-old DJ-1 WT and DJ-1 KO mice. D) Quantitative analysis and graphic representation of PGC1α signal intensity in the retina lysates described above. E) Representative western blot for POLG and β-actin in retina lysates from 3- and 15-month-old DJ-1 WT and DJ-1 KO mice. F) Quantitative analysis and graphic representation of POLG signal intensity in the retina lysates described above. β-actin was used as a housekeeping normalizer, and relative protein expression was calculated by using 3-month-old WT as a normalizer. Data are represented as mean ± SEM; n=3-4 mice per group; statistical analysis using unpaired Student’s t test; significance: p≤0.05 depicted in the figure.

### 2.4 Mitochondrial DNA lesions are not induced by the absence of DJ-1 in RPE and retina

The mtDNA is a circular DNA molecule containing 16,569 DNA base pairs encoding 37 genes, 13 of which encode for polypeptides required for the electron transport chain (ETC), including seven genes encoding for subunits within CI (*MT-ND1*, *MT-ND2*, *MT-ND3*, *MT-ND4*, *MT-ND4L*, *MT-ND5*, *MT-ND6*), one for CIII (*MT-CYB*), three for CIV (*MT-CO1*, *MT-CO2*, *MT-CO3*) and two for CV (*MT-ATP6*, *MT-ATP8*), in addition to two ribosomal RNAs (*MT-RNR1* and *MT-RNR2*) and 22 transfer RNAs (41) (Fig. 6A). The mitochondrial cascade hypothesis proposes that, with age, modifications and mutations in mtDNA accumulate in somatic brain cells, influencing disease phenotypes (42). Moreover, a previous study had identified increased mtDNA damage in the RPE of patients with AMD progression (10). Thus, we performed long extension PCR (LX-PCR) for the ∼16.3 kb of mtDNA by discretely dividing the mtDNA sequence into four regions, RI-RIV and calculating the frequency of mtDNA lesions/10kb (Figs. 6A to 6C). Our analysis in the RPE detected a trend of increase in the accumulation of mtDNA lesions in 15-month-old DJ-1 WT and DJ-1 KO RPE in all regions compared to 3-month-old mice (Fig. 6B). In the retina, we detected a trend of increase in the accumulation of mtDNA lesions/10kb in the 3-month-old DJ-1 KO in all regions compared to 3-month-old DJ-1 WT, and in the 3- and 15-month-old DJ-1 WT mice in RI, RII, and RIII. A significant increase in mtDNA lesions/10kb accumulation in the 15-month-old DJ-1 WT RIV was detected compared to 3-month-old DJ-1 WT (Fig. 6C). We further our analysis by performing deep RIV sequencing for age groups and genotypes using the Oxford nanopore platform. In the RPE, the RIV region sequence analyses suggested no significant changes in the prevalence of mismatch, deletion, insertion, and 6mA methylation between the 3- and 15-month-old DJ-1 WT and DJ-1 KO mice. We also detected an increased frequency of mismatches over deletions, insertions, and mtDNA N6-methyladenine (6mA) methylations in the 3- and 15-month-old DJ-1 WT and DJ-1 KO mice retinas (Suppl. Fig. 2A). Similarly, an increased frequency of mismatches over deletions, insertions, and 6mA methylations in the 3- and 15-month-old DJ-1 WT and DJ-1 KO mice retinas was detected (Suppl. Fig. 2B). Overall, deep sequencing data suggested that mismatch, a form of lesion, is the most abundant event in the RIV of mitochondrial DNA in both RPE and retina, independent of aging and genotype.

**Figure 6.**
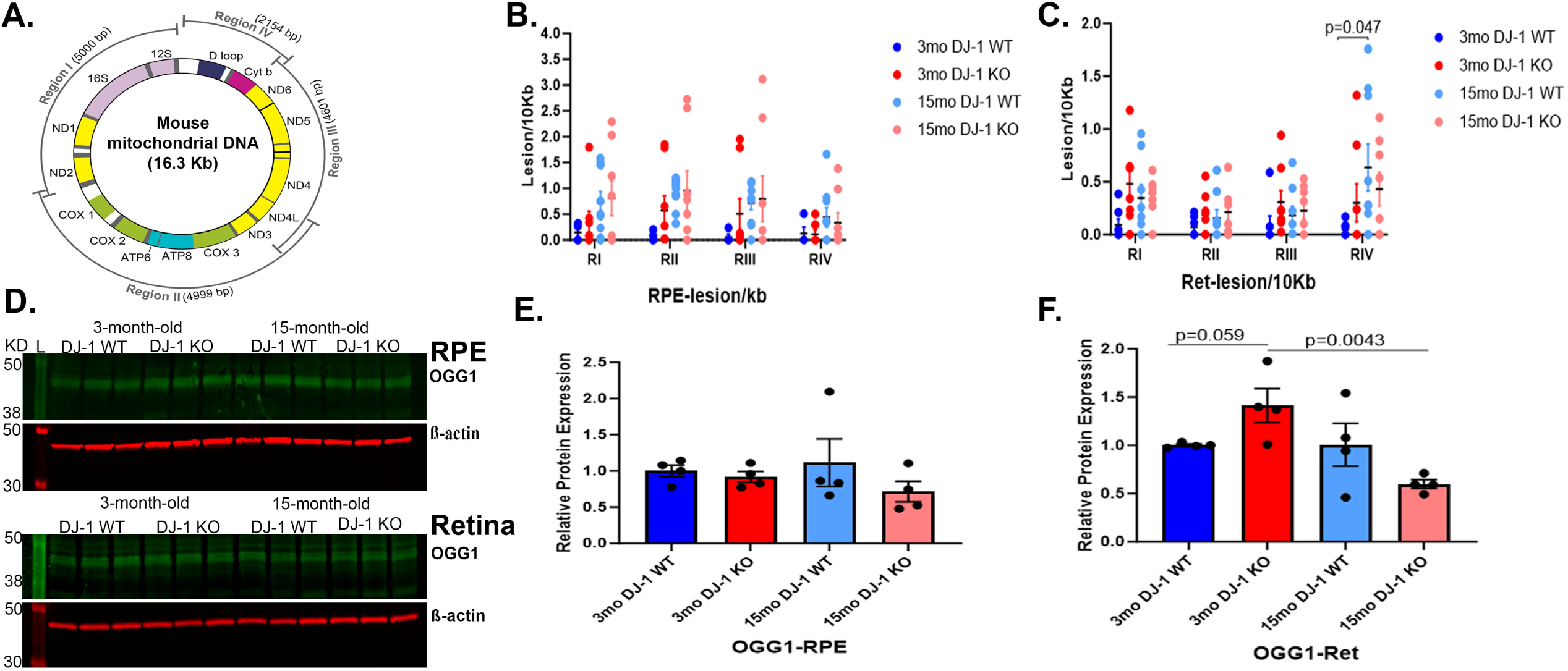
Effects of aging and DJ-1 loss in mtDNA lesion accumulation and mtDNA repair. A) 15ng of genomic DNA from RPE and retina was used for long extension PCR (LX-PCR) using primers designed to amplify specific long base pair regions of the mitochondrial genome Regions (R) I-IV. Quantitative analysis and graphic representation of lesion/10kb in RPE B) and C) retina of 3-month-old DJ-1 WT (dark blue dots), 3-month-old DJ-1 KO (dark red dots), 15-month-old DJ-1 WT (light blue dots), and 15-month-old DJ-1 KO (light red dots). D) Representative western blot for OGG1 and β-actin in RPE and retina. D) Lysates from 3- and 15-month-old DJ-1 WT and DJ-1 KO mice described above. E, F) Quantitative analysis and graphic representation of OGG1 signal intensity in the RPE and retina, respectively. Data for lesion/10kb are represented as mean ± SEM; n=3-4 mice per group; Two-way Anova with Tukey’s multiple comparisons with significant changes are depicted. Data for OGG1 are represented as mean ± SEM; n=3-4 mice per group; β-actin was used as housekeeping normalizer, and relative protein expression was calculated by using 3-month-old DJ-1 WT as normalizer; statistical analysis using unpaired Student’s t test; significance: p≤0.05 depicted in the figure.

In the face of these results, we evaluated the levels of enzyme 8-oxo guanine DNA glycosylase 1 (OGG1) involved in base excision repair. Western blot analyses of OGG1 in the RPE and retina were carried out (Fig 6D), and signal intensities were calculated. In RPE, OGG1 levels were comparable between 3-month-old DJ-1 WT and DJ-1 KO mice. With aging, we observed a trend of decrease in OGG1 levels in 15-month-old DJ-1 KO compared to 15-month-old DJ-1 WT (Figs. 6D, 6E). In the retina, the OGG1 levels were significantly increased in the 3-month-old DJ-1 KO compared to 3-month-old DJ-1WT mice. With aging, we observed a trend of decrease in OGG1 levels in 15-month-old DJ-1 KO compared to 15-month-old DJ-1 WT (Figs. 6D, 6F). Overall, increased base-excision repair enzyme OGG1 suggests enhanced DNA repair in the 3-month-old DJ-1 KO retina.

### 2.5 Antioxidant protein SOD2 levels diminish in the retina with DJ-1 loss in aging

Mitochondrial stress triggered by excess ROS generated by mitochondrial enzymes rich in various oxidoreductases and impairment of defensive antioxidant systems are thought to be contributors to mitochondrial dysfunction (43). Western blots analyzed the levels of mitochondrial superoxide dismutase (SOD2), an enzyme localized in the mitochondria that protects cells from damage caused by ROS. In the RPE, the SOD2 levels in 3-month-old DJ-1 WT and DJ-1 KO were comparable. With aging, no statistically significant differences were observed between the 3-month-old and 15-month-old groups of mice, but 15-month-old DJ-1 KO mice showed a trend of decrease in SOD2 levels when compared to 15-month-old DJ-1 WT mice (Fig 7A, 7B). In the retina, SOD2 levels showed an increased trend in 3-month-old DJ-1 KO compared to 3-month-old DJ-1 WT. With aging, the 15-month-old DJ-1 KO showed a significant decrease in SOD2 compared to 15-month-old DJ-1 WT mice (Fig 7C, 7D). Overall, the SOD2 decrease in aged DJ-1 KO retinas suggests an increase in mitochondrial stress in the retinas of 15-month-old DJ-1 KO mice, which may contribute to the mitochondrial changes detected in our study.

**Figure 7.**
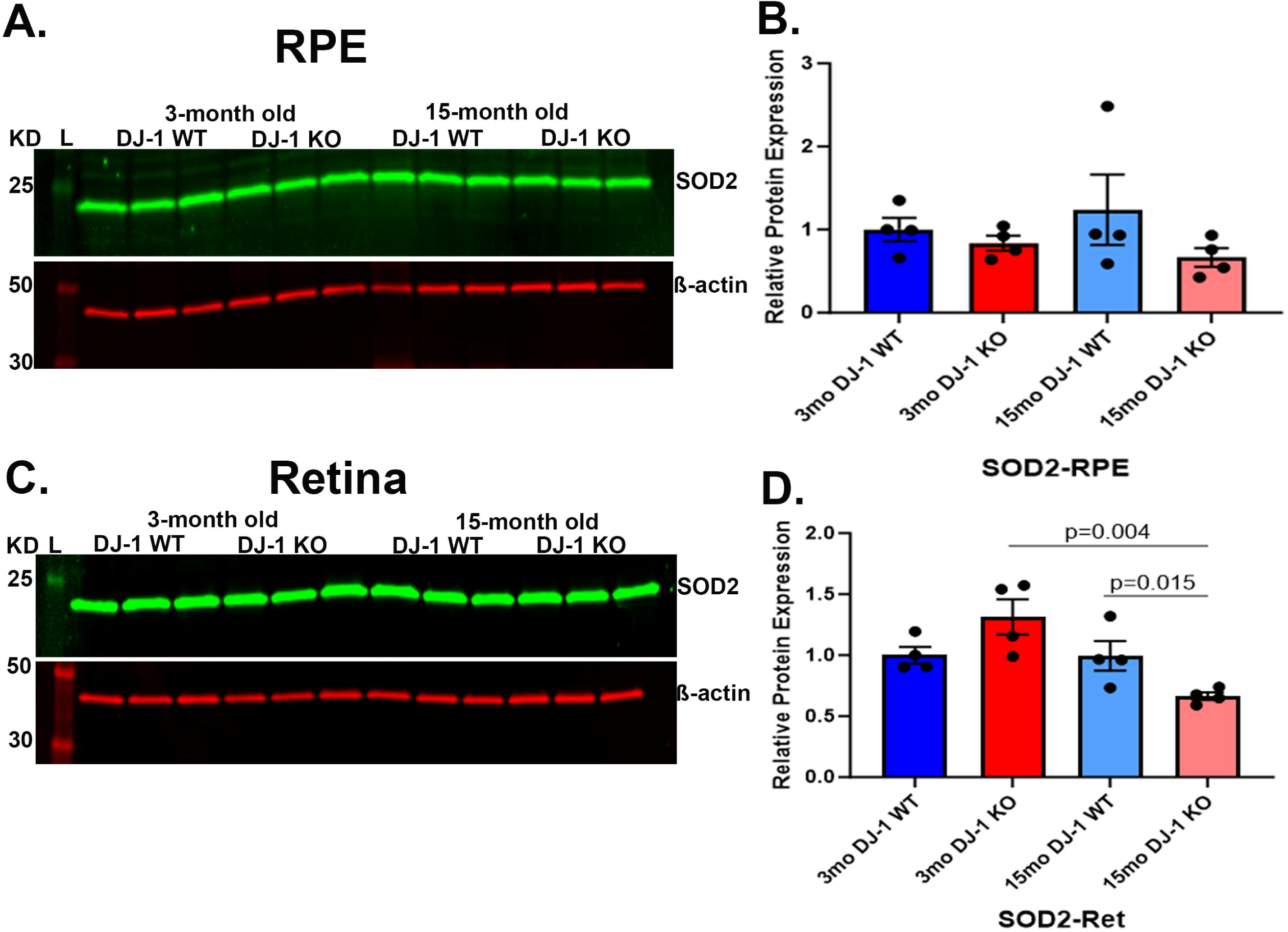
Effects of aging and DJ-1 loss in mitochondrial stress. A) Representative western blot for SOD2 and β-actin in RPE lysate from 3- and 15-month-old DJ-1 WT and DJ-1 KO mice. B) Quantitative analysis and graphic representation of SOD2 signal intensity in RPE. C) Representative western blot for SOD2 and β-actin in retina lysates from 3- and 15-month-old DJ-1 WT and DJ-1 KO mice. D) Quantitative analysis and graphic representation of SOD2 signal intensity in the retina. β-actin was used as a housekeeping normalizer, and relative protein expression was calculated using 3-month-old DJ-1 WT as normalizer. Data are represented as mean ± SEM; n=3-4 mice per group; statistical analysis using unpaired Student’s t test; significance: p≤0.05 depicted in the figure.

## 3. DISCUSSION

During aging, the retina and RPE undergo metabolic dysregulation and mitochondrial dysfunction accompanying deteriorating visual function (44–47). Mitochondrial dysfunction is also associated with several retinal neurodegenerative diseases, including AMD, diabetic retinopathy (DR), and glaucoma (6, 45, 48). In these diseases, a major cause of mitochondrial dysfunction is oxidative stress, which is characterized by increased ROS and decreased antioxidant defense system activity (49–52). DJ-1, also known as PARK7, is a protein renowned for protecting neurons from oxidative stress (22–24).

The DJ-1 protein plays a pivotal role in regulating mitochondria’s response to oxidative stress by translocating to the mitochondria, which aids in the clearance of generated ROS (25, 26). Studies from our group have established increased expression of DJ-1 in RPE in vitro and RPE from AMD human donors as a response to increased levels of oxidative stress (29). Further, we have also reported that loss of DJ-1 accelerates aging in the retina and RPE in DJ-1 KO mice (31) and makes them more susceptible to oxidative damage even under low oxidative stress independent of age (32). Moreover, we have also shown that, in vitro, ARPE-19 and primary human fetal RPE monolayers overexpressing several DJ-1 constructs displayed morphological changes in the mitochondrial structure and mitochondrial membrane potential, suggesting a potential role of DJ-1 in mitochondrial function homeostasis (53). This study comprehensively examined the mitochondria of the retina and RPE of the DJ-1 KO mouse during aging. Our data indicated that the DJ-1 KO retinas and RPE display signs of morphological abnormalities and physiological dysfunction associated with mt dysfunction.

Our ex vivo Seahorse analysis on freshly dissected ex vivo retinal tissues suggested DJ-1 loss might lead to energy deficiency driven by reduced mitochondrial respiration and glycolysis in the retina that can be detected at 3-months of age. Previous studies established that DJ-1 maintains mitochondrial homeostasis by regulating mitochondrial CI (28, 54). DJ-1 is found in the mitochondrial matrix and intermembrane space, where it binds to mitochondrial CI subunits and controls its assembly and activity. Specifically, depletion of DJ-1 significantly reduced the OCR, ATP production (55), and CI activity (28).

We determined that DJ-1 deficiency can negatively alter OXPHOS subunit expression in RPE and retina at baseline levels and with aging. In young mice, the ATP5A subunit of OXPHOS CV is significantly decreased in RPE, while OXPHOS CI and CIV were significantly decreased in the retina. ATP5A is a nucleus-encoded peptide of the F1subunit of ATP synthase involved in the catalytic production of ATP by the OXPHOS system. This observation suggests that DJ-1 loss can be associated with lower ATP production and, therefore, mitochondrial dysfunction in RPE. It also corroborates a previous study reporting a decrease in ATP synthase B subunit (CV) protein expression and a decrease in ATP synthase enzymatic rate in isolated neurons and HEK-293T cells lacking DJ-1 function (56). Nevertheless, a significant decrease of NDUFB8 (CI), SDHB (CII), and MTCO1 (CIV) in RPE of 15-month-old DJ-1 KO in comparison to 15-month-old DJ-1 WT implicates the presence of DJ-1 being essential for retaining their expression with aging.

In the retina of 3-month-old DJ-1 KO mice, there is a significant downregulation of the protein expression of NDUFB8 (CI) and MTCO1 (CIV) compared to 3-month-old WT mice. It is known that CIV loss triggers oxidative damage and partial degradation of critical C I subunits (57, 58). Therefore, it is possible that CIV loss would be an important contributing factor in CI deficiency in DJ-1 KO retina. The significant decrease in MTCO1 (CIV) in the retina of 15-month-old DJ-1 WT and DJ-1 KO mice compared to their 3-month-old counterparts suggests an aging hallmark. Indeed, several studies have shown age-related changes in protein expression of CIV related to mitochondrial dysfunction (59, 60). Finally, contradictory changes were reported in regulating CIV by DJ-1 in different cells. Isolated mitochondria from primary neurons and in the forebrain of mice lacking DJ-1 displayed lower free *versus* supercomplexes-assembled CI and CIII ratios, and the CIV loss in the samples (61). However, an increase of CIV was reported in DJ-1 null dopaminergic neuronal cell line lysates (55).

A noteworthy aspect of mitochondrial dysfunction is changes in mtDNA content. MtDNA content, a measure of mitochondrial abundance, has been identified to be altered with age and disease in humans (62). Recent studies suggest that in several human genetic mitochondrial disorders, mitochondrial copy number is increased (63), whereas it tends to decrease in certain neurodegenerative diseases (64, 65). Our assessment of mtDNA content using mtco1 and 16s suggested a two- and three-fold increase in RPE of 3- and 15-month-old DJ-1 KO mice compared to age-matched DJ-1 WT. A previous study detected increased mtDNA copy number with age in the retina of C57BL male mice, analyzing the Cox2 gene (66). Increasing mtDNA content in young and old RPE and old retina in DJ-1 KO mice could be a compensatory measure to maintain the OXPHOS activity, as mtDNA expression is a prerequisite for the biogenesis of the OXPHOS system.

To further understand the increased mtDNA content, we analyzed protein expression of PGC1α and POLG aiming to assess mitochondrial biogenesis and mtDNA replication. Downregulation of PGC1α can result in reduced mitochondrial antioxidant capacity and heightened ROS production (67). Thus, the detected decrease of PGC1α levels in RPE but not retina of 3- and 15-month-old DJ-1 KO compared to age-matched DJ-1 WT suggests increased stress in the RPE. The decrease of PGC1α levels within groups during aging agrees with many studies which suggest lower PGC1α with aging in several tissues (38, 68, 69). Interestingly, the PGC1α levels in the retina decreased only in DJ-1 WT mice during aging.

Analyses of RPE detected decreased levels of POLG between 3- and 15-month-old DJ-1 KO. In the retina, POLG levels increased in 15-month-old DJ-1 KO compared to 15-month-old DJ-1 WT. Because the integrity and normal replication of mtDNA are crucial to the health of mitochondria (70), our results suggest decreased health of the DJ-1 KO RPE mitochondria compared to the retinal mitochondrial health. POLG activity and mtDNA copy number can also be regulated by POLG methylation in a tissue-specific manner (71, 72). Further analysis of POLG methylation is needed to understand the role of POLG in the retina and RPE of our samples.

We also investigated the changes in levels of another enzyme associated with DNA repair functions in the retina and RPE, OGG1. We observed a trend of decrease in RPE OGG1 levels in 15-month-old DJ-1 KO compared to 15-month-old DJ-1 WT. In the retina, the OGG1 levels were significantly increased in the 3-month-old DJ-1 KO compared to 3-month-old DJ-1WT mice. Previous studies reported increased mtDNA damage and down-regulation of OGG1 in aged mice RPE and choroid (73). The level of OGG1 has been reported to be approximately two-fold higher in young macular primary RPE cells compared with aged macular primary RPE cells. Furthermore, the level of OGG1 decreased by almost half in AMD macular RPE cells compared with aged macular RPE cells (74, 75). OGG1 is also considered an oxidative stress marker protein (75, 76). Lastly, we analyzed the mitochondrial redox status by quantifying mitochondrial SOD2 and levels in RPE and retina. In the retina, SOD2 levels showed an increased trend in 3-month-old DJ-1 KO compared to 3-month-old DJ-1 WT. With aging, the 15-month-old DJ-1 KO showed a significant decrease in SOD2 compared to 15-month-old DJ-1 WT mice. Previous studies reported a direct correlation between levels of DJ-1 SOD2 (77–79), suggesting that the DJ-1 function may be uniquely regulated in the retina and RPE. Overall, the increase observed in OGG1 and SOD2 in the DJ-1 KO retinas of 3-month-old mice may indicate increased oxidative stress in the mitochondria of these mice at young ages.

Because mtDNA integrity is related to mitochondrial function, we also performed LX-PCR to score disruption of the PCR reaction, represented by the number of lesions per 10 kb of mtDNA in the complete mouse mtDNA. This assay detected a significant increase in mtDNA lesions/10kb accumulation in the 15-month-old DJ-1 WT RIV compared to the 3-month-old DJ-1 WT. Deep sequencing of the RIV region, which contained the sequence for cytb and D-loop, also suggested no significant differences in % frequency/base for mismatch, deletion, insertion, and 6mA methylation in RPE and retina with loss of DJ-1 and aging. Our in-depth sequencing data for RIV determined that the % frequency of mismatch was higher than deletion, insertion, and 6mA methylation and implied that mismatch was the more favored in mtDNA. The significance of these sequence changes needs to be further explored. However, it has been reported that mtDNA 6mA could repress DNA binding and bending by mitochondrial transcription factor (TFAM) and that levels of 6mA in mtDNA could be further elevated in conditions of stress and disease resulting potentially in mitochondrial dysfunction (80, 81).

In conclusion, this study provides a detailed analysis of potential processes linking DJ-1 and aging as dual drivers of mitochondrial dysfunction in RPE and retina. The data from this study can be relevant to retinal degenerative diseases like AMD, whose pathogenesis has been linked to aging, chronic oxidative stress, and mitochondrial dysfunction.

## 4. MATERIALS AND METHODS

### 4.1 Mice

All animal procedures were approved by the Institutional Animal Care and Use Committee (ARC 00002191) at the Cleveland Clinic. Mice were housed in individually ventilated cages for 14 hours of light and 10hours of darkness cycles and were provided regular chow ad libitum. The experiments were performed in male and female 3-month-old (young) and 15-month-old (Old) DJ-1 KO and DJ-1 WT mice. Mice were genotyped as previously described previously (30).

### 4.2 Protein extraction and Western blotting

Retinas were mechanically detached from the RPE/choroid under a dissecting microscope in Hanks’ Balanced Salt Solution without Ca2+, Mg2+ solution. Isolated retinas and RPE were lysed in RIPA buffer (cat#J63324, Thermo Scientific, Waltham, MA, USA) containing protease inhibitors and phosphate inhibitors (Cat#P8340, P5626, P0044, Sigma-Aldrich, MO, USA). Retinas were sonicated twice for 15 seconds, and RPE was lysed by passing through a 27 1/2 G syringe needle, followed by incubation on ice and vortexed every 5 minutes for 20 min after. After lysis, retina and RPE lysates were centrifuged at 4 °C for 10 minutes at 14000 rpm. The supernatants were collected, and the protein concentration was quantified using a Micro BCA Kit (Cat#23235, Thermo Scientific, WA, MA, USA). 30µg of protein for retina and RPE lysate were resolved on 4-20% Tris-Glycine SDS PAGE (cat# 5671093, Bio-Rad, Hercules, CA, USA) for probing of OXPHOS, PGC1α, Polg and SOD2; and on 12% Tris-Glycine SDS PAGE (Cat#5671043, BioRad, Hercules, CA, USA) for probing of OGG1. Gels were transferred on PVDF membrane (Immobilon-FL, Cat#IPFL00010, Merck Millipore Ltd., MA, USA) under wet conditions at constant volts for 1.5 hours. Membranes were incubated with the following antibodies: Total OXPHOs cocktail (Cat#ab110413, Abcam, Cambridge, UK), SOD2 (Cat#ab13533, Abcam, Cambridge, UK), Polg (Cat#A1323, ABclonal, Woburn, MA, USA), PGC1α (Cat#NBP1-04676, Novus Biologicals LLC, Centennial, CO, USA), OGG1 (Cat#15125-1-AP, Proteintech, Rosemont, Illinois, USA); Gapdh (Cat310494-1-AP, Proteintech, Rosemont, Illinois, USA) or β-actin (Cat#8H10D10, Cell Signaling Technologies, Danvers, MA, USA) was used as loading control. The following secondary antibodies provided by Licor Biosciences (Lincoln, NE, USA) were used: Anti-mouse IRDye®680RD, Anti-rabbit IRDye®680RD and anti-mouse IRDye®800CW. The immunoreactive bands were visualized using Oddessey CLx (Licor Biosciences, NE, USA). The bands were quantified in Licor’s ImageStudio V5.2 software. Fold changes were calculated using the 3-month-old DJ-1 WT signal intensities as a normalizer.

### 4.3 DNA extraction and mtDNA content

The retina was mechanically detached from RPE, as described above. Lysis of isolated tissue was performed in an ATL buffer containing Proteinase K, which is available from the DNeasy blood and tissue kit (Cat#69506, Qiagen GmbH, Germany). Total genomic DNA (gDNA) was isolated using DNeasy blood and tissue kit (Cat#69506, Qiagen GmbH, Germany) according to manufacturer’s instructions. 15ng of genomic DNA from RPE and retina was used as a template. qPCR was performed on Quantstudio 3 real-time PCR system (Life Technologies, Carlsbad, CA, USA) using ABI Powerup Syber green mastermix (cat#A25742, Thermo Scientific, Waltham, MA, USA). The mtDNA genes mtco1 and 16s, as well as the nuclear gene B-globin, were amplified. The PCR primer sequences are provided in Table 2. Primer concentrations for mtco1 (2 mM), 16s (5 mM), and b-globin (10 mM) were optimized using the standard curve method. The qPCR was performed for 40 cycles at 95^0^C for 15 seconds, 57^0^C for 20 seconds and 72^0^C for 30 seconds.Relative mtDNA content was determined as previously described (82). Obtained Ct values were normalized to 3-month-old DJ-1 WT mice.

### 4.4 Long-extension PCR (LX-PCR) and Qubit quantification

Total genomic DNA was used to amplify mtDNA. The 16.3 kb mouse mtDNA sequence was divided into four discrete regions-RI-RIV (Table 1). The sequence for forward and reverse primers for RI and RII were procured from Oligotech services (Merck, Darmstadt, Germany), and forward and reverse primers for RIII and RIV were designed in the lab using http://basic.northwestern.edu/biotools/OligoCalc.html tool (83). After empirical optimization, a 15ng gDNA template and 10 mM concentration of forward and reverse primer were used to amplify gDNA for each region (RI-RIV). A half-template control and no-template control reaction tube were always run to verify linearity and negative control, respectively. All regions were amplified using the following PCR cycling parameters: Initial denaturation at 94^0^C for 3 minutes, followed by 19 cycles for RI&RIV and 18 cycles for RII& RIII of denaturation at 94^0^C for 15 seconds, annealing at 50^0^C for 20 seconds, extension at 68^0^C for 5 minutes and final extension at 68^0^C for 10 minutes. A small 16s ∼200bp fragment was amplified to correct for variations in mtDNA copy numbers across the samples. The PCR products were sequenced to verify the correct amplicon. All the PCR products were quantified using Qubit 4 (Thermo Scientific, Waltham, MA, USA) according to the manufacturer’s instructions. Lesion/10Kb was determined as previously described (84).

**Table1.**
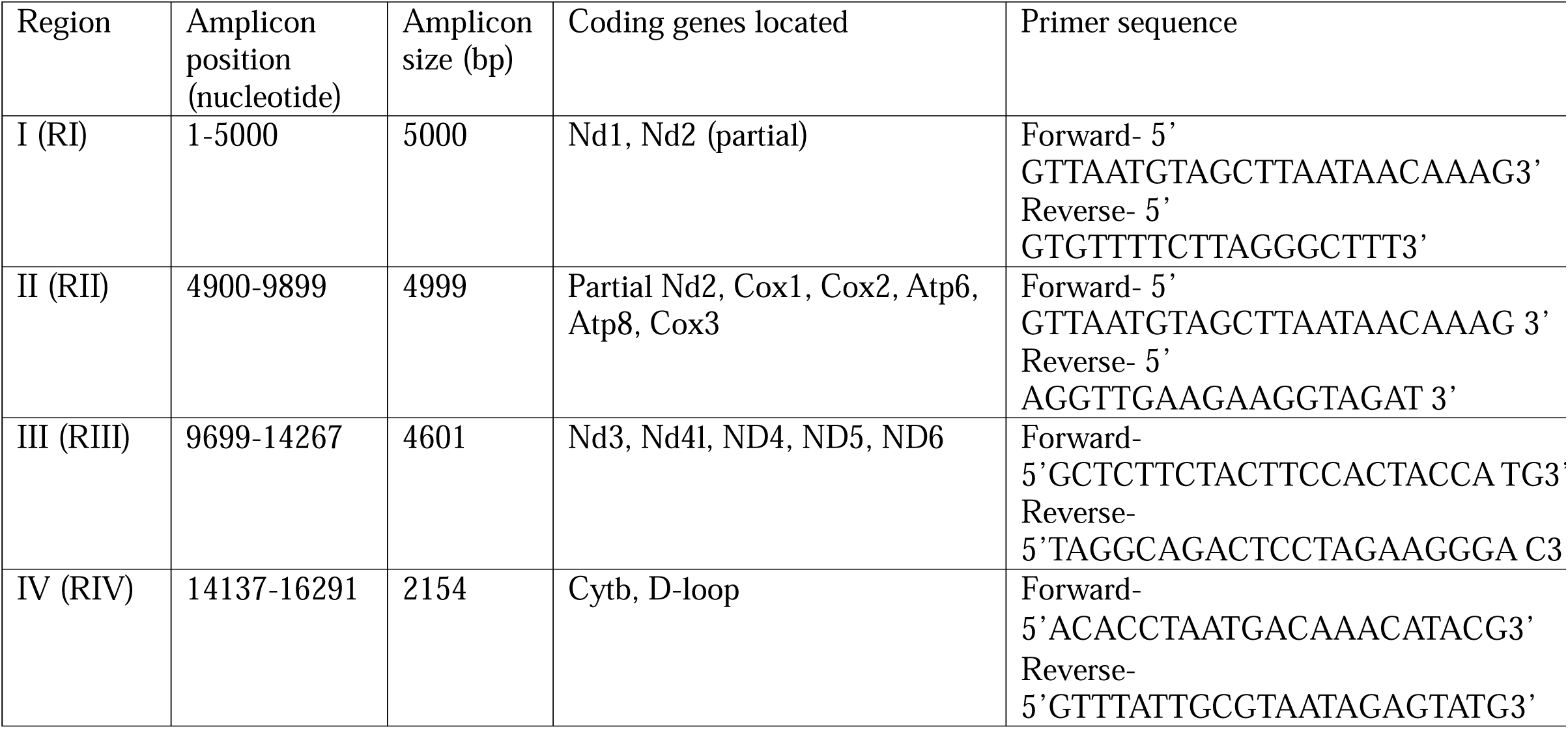
Details of regions of mouse mitochondrial DNA sequence amplified by LX-PCR.

**Table 2.**
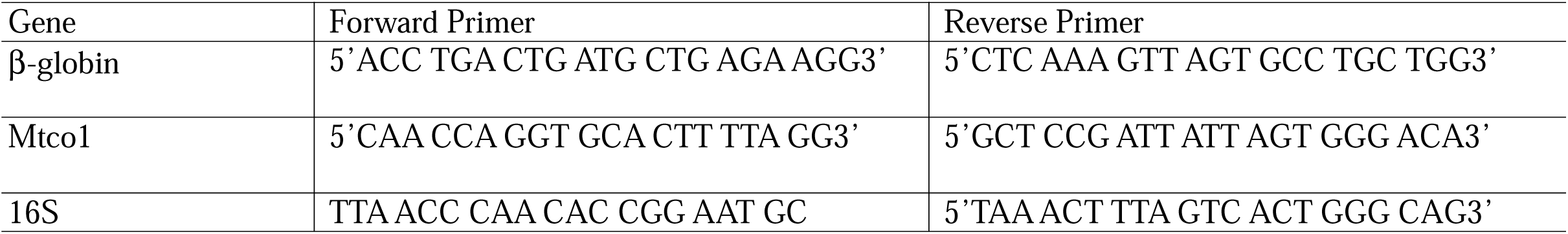
Primer sequences for relative mtDNA content assay.

### 4.5 Deep sequencing for RIV amplicons

mtDNA RIV region was amplified as described above. PCR products were purified using the QIAquick PCR purification kit (cat#28104, Qiagen, GmbH, Germany) as per the manufacturer’s instructions. The purity and integrity of the purified product were checked using Qubit4 and Nanodrop (Thermo Scientific, Waltham, MA, USA). Purified PCR product at 50ng/ul concentration was sent to Plamidsaurus Inc. (Louisville, KY, USA) for sequencing using Oxford nanopore technology.

### 4.6 Seahorse assay

Seahorse analysis was conducted on freshly dissected ex vivo retinal punches to determine oxygen consumption rate (OCR) and extracellular acidification rate (ECAR) using the Seahorse XFe24 Islet Capture FluxPak (Agilent, 103518-100), as previously described (33). One day prior to the experiment, sensor cartridges were soaked in calibration solution overnight at RT. The Seahorse DMEM assay medium containing 6 mM glucose, 0.12 mM of pyruvate, and 0.5 mM of glutamine was prepared fresh the day of the experiment. Mesh inserts in the islet capture microplate were coated with a cell attachment medium (Cat# 354240, Corning Cell-Tak, Corning, NY,USA) to improve tissue attachment. Retinas were dissected in assay medium, and a 1mm diameter biopsy puncher was then used to take five separate retinal punches from each retina. Punches were then placed on mesh inserts and loaded into the islet capture microplate containing 500 μL assay medium. For mitochondria stress assay, punches were incubated with 5 μM of Bam15 (Cat# ST056388, TimTec LLC, Tampa, FL, USA) and 1 μM of each rotenone (Cat# R8875, Sigma-Aldrich, St. Louis, MO, USA)/antimycin A (Cat# R8674, Sigma-Aldrich, St. Louis, MO, USA). For the glycolytic stress assay, punches were incubated with 1 μM mix of rotenone/antimycin A mix and 50 mM 2-DG (Cat# D6134, Sigma-Aldrich, St. Louis, MO, USA). The assay program in the analyzer was then prepared as follows: 5 cycles of measurements for baseline, injected the first drug, followed by 4 cycles of measurements, then injected the second drug, followed by another 4 cycles of measurements. Each cycle was composed of mix (3 min), wait (2 min) and measure (3 min). At the completion of the assay, data was collected, and statistical analysis was performed in Graphpad Prism 9 using one-way ANOVA, p-value < 0.05.

### 4.7 Statistics

All data were analyzed using GraphPad Prism v10.0 (GraphPad Software, La Jolla, CA). Seahorse data are presented as mean± SD and statistically analyzed using One-way Anova. All other data are presented as mean± SEM. Groups for western blot were compared by unpaired t-test. Two-way Anova compared groups for lesion load and mutations with Tukey’s correction.

## ACKNOWLEDGMENTS

The authors thank Maximilian Campbell and Benjamin A. Routhier for their technical assistance. The authors also thank David Schumick, BS, CMI for preparation of the illustration included in the manuscript.

## Supplemental Figure

**Figure S1. Effects of aging and DJ-1 loss in ECAR measured by Seahorse analysis.** Five measurements were taken before treatment (basal glycolysis), four measurements after 1 μM rotenone/antimycin A injection (maximal glycolysis), and four measurements after 50 mM 2-DG injection (glycolytic reserve capacity). A) Unnormalized ECAR measurements in retinal punches from 3-month-old DJ-1 WT (dark blue), 3-month-old DJ-1 KO (dark red), 15-month-old DJ-1 WT (light blue), and 15-month-old DJ-1 KO (light red) mice. B) Basal ECAR, C) Maximal ECAR, and D) Glycolytic reserve capacity (GRC) of mice described above. Error bar indicates SEM. Data points = technical replicates / individual retina punches from n = 3-5 mice. Statistics: one-way ANOVA; significance: p≤0.05 depicted in the figure.

**Figure S2. Effects of aging and DJ-1 loss in mtDNA sequencing corresponding to cytb and D-loop (RIV) using nanopore sequencing.** Quantification of % frequency for mismatch, deletion, insertion, and 6mA methylation in the A) RPE and B) from 3-month-old DJ-1 WT (dark blue circles), 3-month-old DJ-1 KO (dark red circles), 15-month-old DJ-1 WT (light blue triangles), and 15-month-old DJ-1 KO (light red triangles) mice. Data are represented as mean ± SEM; n=3-4 mice per group. Statistics: two-way ANOVA with Tukey’s multiple comparisons; significance: p≤0.05 depicted in the figure.

