## Supplementary figures and images for "DJ-1 deficiency and aging: dual drivers of retinal mitochondrial dysfunction"

### Supplemental Figures 1 and 2

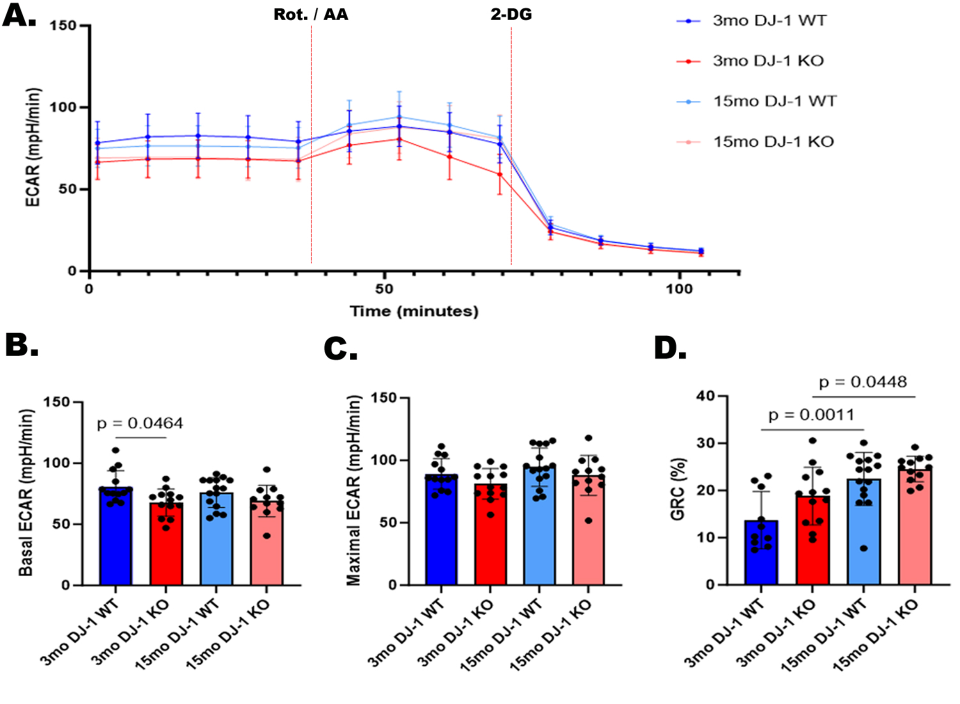

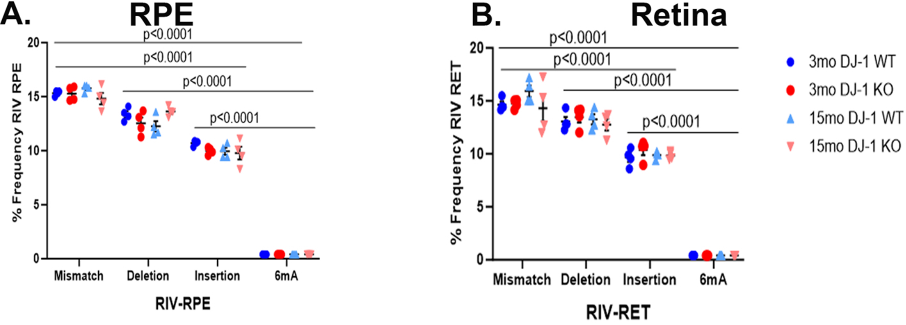
